# DJ-1 depletion slows down immunoaging in T-cell compartments

**DOI:** 10.1101/2021.05.21.445139

**Authors:** Ni Zeng, Christophe Capelle, Alexandre Baron, Severine Cire, Cathy Leonard, Djalil Coowar, Haruhiko Koseki, Astrid M. Westendorf, Jan Buer, Dirk Brenner, Rejko Krüger, Rudi Balling, Markus Ollert, Feng Q. Hefeng

## Abstract

Decline in immune function during aging increases susceptibility to different aging related diseases. However, the underlying molecular mechanisms, especially the genetic factors contributing to imbalance of naïve/memory T-cell subpopulations, still remain largely elusive. Here we show that loss of DJ-1 encoded by *PARK7*/DJ-1, causing early-onset familial Parkinson’s disease (PD), unexpectedly delayed immunoaging in both human and mice. Compared with two gender-matched unaffected sibling carriers of similar ages, the index PD patient with DJ-1 deficiency showed a decline in many critical immunoaging features, including almost doubled frequencies of non-senescent T cells. The observation of a ‘younger’ immune system in the index patient was further consolidated by the results in aged DJ-1 knockout mice. Our data from bone marrow chimera models and adoptive transfer experiments demonstrated that DJ-1 regulates several immunoaging features via hematopoietic-intrinsic and naïve-CD8-intrinsic mechanisms. Our finding suggests an unrecognized critical role of DJ-1 in regulating immunoaging, discovering a potent target to interfere with immunoaging- and aging-associated diseases.

## Main text

Decline in immune function during aging, known as immunoaging or immunosenescence (Gruver et al. 2007, Nikolich-Žugich 2018), increases susceptibility to different complex non-communicable (Niccoli and Partridge 2012) and infectious diseases (CDC 2019, Akbar and Gilroy 2020), therefore creating a primary healthcare burden with the increasing elderly populations worldwide (Aw et al. 2007, WHO 2011). A growing body of evidence has linked the chronic infection, especially cytomegalovirus (CMV) (Nikolich-Zugich 2008, Brunner et al. 2011, Pawelec and Derhovanessian 2011, Fulop et al. 2013) to the cause of immunoaging in the organismal level. A number of pathways (Cavanagh et al. 2011, Mannick et al. 2014, Lanna et al. 2017, Pereira et al. 2020) have been associated with immunoaging. However, the genetic factors contributing to the regulation of the ratio alteration among different T-cell subpopulations during aging as well as the regulation of other immunoaging features still remain largely elusive (Sansoni et al. 2014, Goronzy and Weyand 2019).

Our recent work have demonstrated that DJ-1 depletion (Bonifati et al. 2003) reduced CD4 regulatory T cells (Treg) cellularity only in aged, but not young adult mice (Danileviciute et al. 2019). Since Treg frequency increases during natural aging (Raynor et al. 2012), we hypothesized that DJ-1 might regulate immunoaging in other T-cell compartments. To leverage the translational potential of this work, we started our analysis from an index PD patient carrying the homozygous c.192G>C mutation in the DJ-1 gene and two of his siblings, who are unaffected heterozygous carriers of the same PD causing mutation (Burbulla et al. 2017). In comparison with the unaffected age-matched siblings (**Fig. 1a**, all with the same gender, 56-63-year old at the time of sampling), the patient without DJ-1 expression had an approximately twofold higher frequency of circulating non-senescent CD8 T cells, e.g., CD27^+^CD28^+^ CD8 T cells (**Fig. 1b**). The frequency of senescent or exhausted T cells in the index patient, such as CD57^+^, PD-1^+^, Eomes^+^ and Tbet^+^ CD8 T cells, was reduced to almost half of the two siblings (**Fig. 1c-f**). A similar effect was observed for CD4 T cells (**Extended Data Fig. 1a-f**). In line with our previous observations about the Treg frequency change in aged whole-body DJ-1 KO mice (Danileviciute et al. 2019), the frequency of FOXP3^+^CD4^+^ Tregs also declined in the affected patient versus the two unaffected siblings (**Extended Data Fig. 1g**).

**Figure 1.**
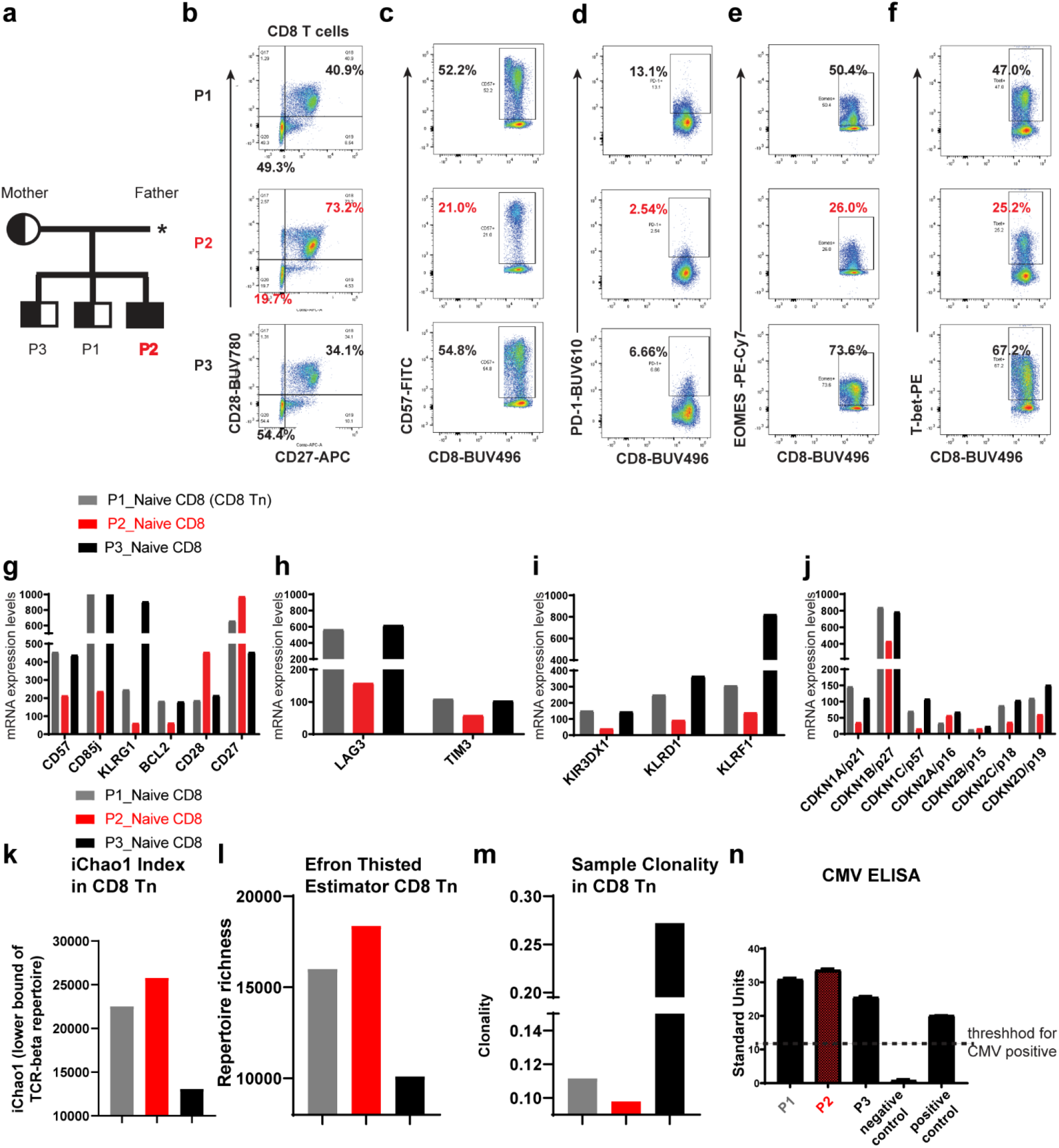
DJ-1 deficiency delays immunoaging in CD8 T cells of an index patient. **a**, Extract of the pedigree of the family carrying the c.192G>C mutation in the DJ-1 gene indicating the three participants [P1 (heterozygous mutation), P2 (homozygous mutation) and P3 (heterozygous)]. The half black square/circle indicates male/female individuals heterozygous for the DJ-1 mutation, the black square indicates the patient carrying the homozygous DJ-1 mutation. *\**Obligate heterozygous mutation carrier although dead at the time of the initial study. **b**, Expression of CD27 and CD28 on peripheral blood CD8 T cells of three participants. **c, d, e, f**, Expression of CD57 (**c**), PD-1 (**d**), EOMES (**e**) and T-bet (**f**) on peripheral blood CD8 T cells of three participants. The enlarged number in the corresponding gate represents the corresponding percentage out of the parent population. **g-j**, mRNA expression of senescence related genes (**g**), exhaustion related genes (**h**), KIR or KLR genes (**i**) and cyclin-dependent kinase inhibitor genes (**j**) in sorted CD8 T cells of the peripheral blood from three participants. **k, l**, Comparison of the lower bound of TCR-beta repertoire (**k**) and richness (**l**) of naïve CD8 T cells among three participants. **m**, The sample clonality index of TCR repertoire of naïve CD8 T cells among three participants. **n**, Serology detection of CMV infection for three participants. The threshold of 11 was decided according to the manufacture instruction.

To more comprehensively characterize the expression of other senescence-related genes, we performed genome-scale transcriptomic analysis in sorted CD8 T cells of the three siblings using microarray. Notably, many immunosenescence marker genes (Goronzy and Weyand 2013), such as killer immunoglobulin-like receptors (KIRs), killer cells lectin-like receptors (KLRs) and exhaustion markers, e.g., KLRG1, CD85j, LAG3, TIM3 and several cyclin-dependent kinase inhibitors including p21, known to increase during aging, have all been decreased substantially in the patient with a homozygous DJ-1 mutation relative to the heterozygous carriers (**Fig. 1g-j**). During aging, the TCR repertoire diversity decreases in naïve T cells (Ahmed et al. 2009, Britanova et al. 2014). In line with other observations, the TCR repertoire diversity, which is positively correlated with iChao1 index (**Fig. 1k**) and richness (**Fig. 1l)**, but negatively correlated with sample clonality index (**Fig. 1m**), was increased in sorted naïve CD8 T cells (CD8 Tn) of the index patient. This also held true for the TCR repertoire diversity in naïve CD4 T cells (CD4 Tn, **Extended Data Fig. 1h-j**). It is well accepted that some chronic infections, especially CMV, markedly accelerate immunoaging(Brunner et al. 2011). We therefore applied serological testing for CMV IgG antibody titers and found that all the three subjects were CMV positive (**Fig. 1n**). Furthermore, there was no clinical evidence for a specific susceptibility to chronic diseases or repetitive infections of all the three siblings. No increase or decrease trend was observed in systematic levels of relevant pro-inflammatory cytokines (Ferrucci and Fabbri 2018) in the index patient versus the two unaffected siblings (e.g., low levels of plasma IFNg, TNFa, IL6 and undetectable or below fit curve range for IL1b, IL4, IL5, IL17a and IL10 among all the three siblings) (**Extended Data Fig. 1k-m**). Thus, these data encouraged us to believe that the reduced immunoaging observed in the index patient was driven by DJ-1 deficiency, but not simply due to CMV infection or other chronic infectious diseases or systematic inflammation from either siblings.

DJ-1 loss-of-function mutations are a rare cause of monogenic PD (Pankratz et al. 2006) and we were unable to identify further patients with *PARK7*-related PD available for biosampling in our extended networks (Boussaad et al. 2020). Therefore, to obtain more statistic power and mechanistic insights, we analyzed whole-body DJ-1 knockout (KO, for simplicity ‘whole-body’ will be left out afterwards unless different lines) mice. Here we examined relevant features of immunoaging in DJ-1 KO mice, which were developed elsewhere to study the impact of DJ-1 on neurodegenerative diseases, i.e., PD (Pham et al. 2010). As a hallmark of natural aging, memory T cells increase while their counterpart, Tn, decrease (Nikolich-Zugich 2008). However, we detected a significantly higher frequency of CD8 naïve T cells (Tn, CD44^low^CD62L^high^) accompanied by a lower frequency of effector memory (CD44^high^CD62L^low^, Tem) in aged (∼45 wks) DJ-1 KO mice versus age-and gender-matched WT (**Fig. 2a-c**). A tendency for reduced frequency of central memory CD8 T cells (CD44^high^CD62L^high^, CD8 Tcm) was also observed in aged DJ-1 KO mice (p=0.07, **Extended Data Fig. 2a**). No change in the memory/naïve compartments of CD8 T cells was observed in young adult (8-15 wks, simplified as ‘young’ later on) DJ-1 KO mice (**Fig. 2a-c, Extended Data Fig. 2a**). Although immunosenescence does not fully overlap with exhaustion(Akbar and Henson 2011), a considerable intersection exists between these two age-related immunophenotypes and their functional consequences. In contrast to the increased PD-1 expression in T cells during natural aging (Lee et al. 2016), we observed a significantly lower expression of the key exhaustion marker PD-1 among homeostatic total CD8 T cells (**Fig. 2d**) and different CD8 subsets (**Extended Data Fig. 2b**) in aged, but not young DJ-1 KO versus WT mice.

**Figure 2.**
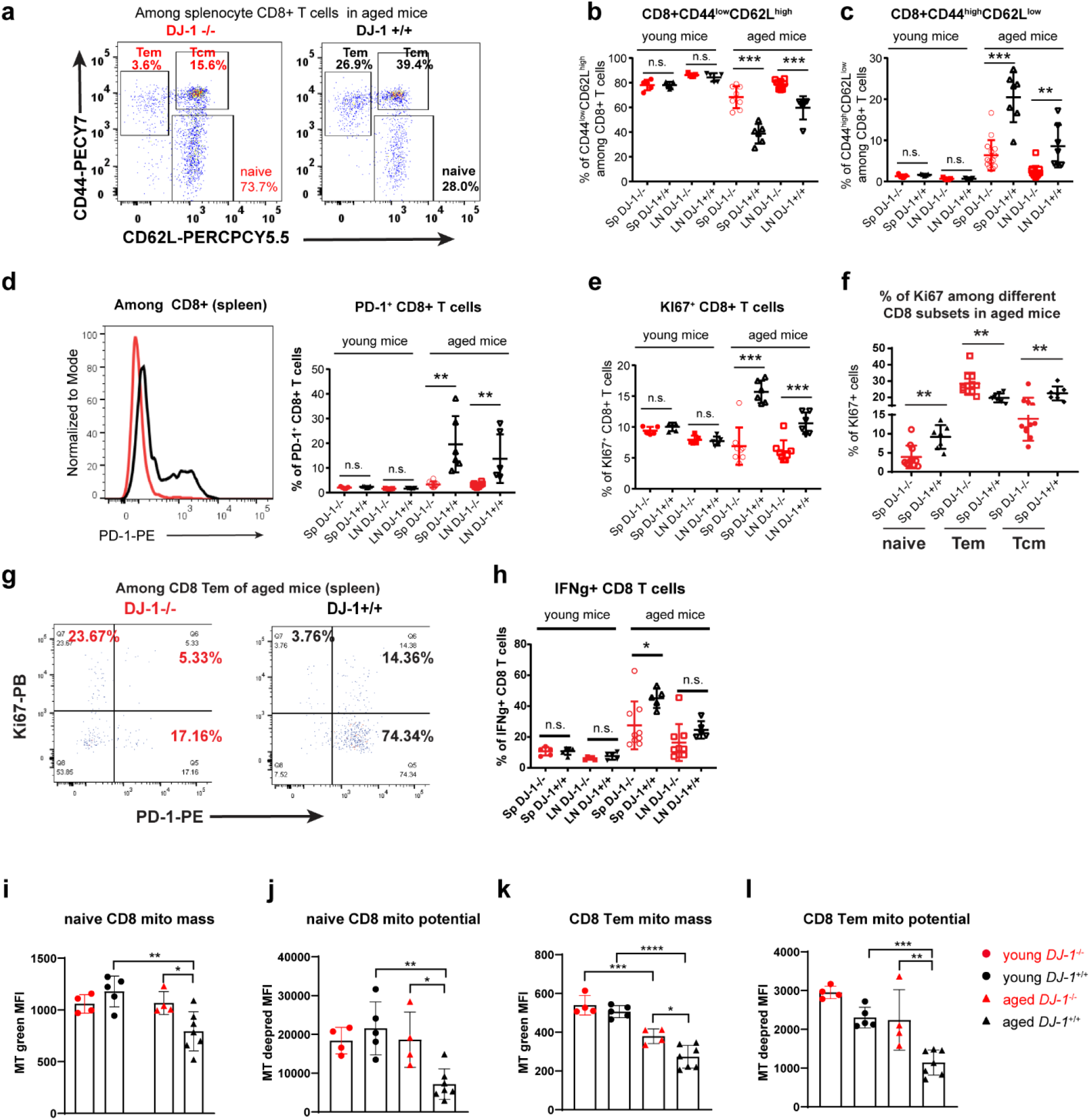
DJ-1 depletion delays immunoaging in murine CD8 T cells. **a**, Representative flow-cytometry plots of CD44 and CD62L expression on total CD8 T cells of aged DJ-1 KO and age and gender-matched WT mice. **b, c**, Percentages of CD44^low^ CD62L^high^ (Tn) (**b**) and CD44^high^ CD62L^low^ (Tem) (**c**) cells among total CD8 T cells of spleen and pLNs from young and aged DJ-1 KO and WT littermates. (young KO, n=5; young WT, n=5; aged KO, n=8; aged WT, n=6; for aged mice, data pooled from 2 independent experiments). **d**, Representative histogram overlay of PD-1 expression among total CD8 T cells in spleen of aged mice (left panel) and percentages of PD-1^+^ cells among total CD8 T cells (right panel). **e**, Percentages of Ki-67^+^ cells among total CD8 T cells. **f**, Percentage of Ki-67^+^ cells among splenic CD8 Tn, Tem and Tcm in aged DJ-1 KO and age-and gender-matched WT mice. **g**, Representative flow-cytometry plots of Ki-67 and PD-1 among splenic CD8 Tem of aged mice. **h**, IFNγ production in CD8 T cells of spleen and pLNs after in vitro stimulation using 50 ng/ml of PMA and 750 ng/ml of ionomycin for 5 h. **i, j, k, l**, Comparison of splenic CD8 Tn mitochondrial mass (mito, Mitotracker green MF, **i**) and mitochondrial potential (MitoTracker Deep Red, **j**) of young and aged DJ-1 KO and WT mice. Comparison of CD8 Tem mitochondrial mass (**k**) and mitochondrial potential (**l**) of young and aged DJ-1 KO and WT mice. SP and LN represents spleen and lymph nodes, respectively. Results represent at least four (**b-h**) or three (**i-l**) independent experiments. Data are mean± s.d. The P-values are determined by a two-tailed non-paired Student’s *t*-test. ns or unlabeled, not significant, *P<=0.05, **P<=0.01 and ***P<=0.001.

This observation indicates that the exhaustion process might slow down in aged DJ-1 KO mice. In addition to exhaustion, PD-1 is also regarded as one of the key activation markers of T cells (Ahn et al. 2018). Therefore, to determine whether our observation is based on either reduced activation in general or cell exhaustion, we also assessed proliferation markers, e.g., Ki-67 in different T cell subsets under homeostatic conditions. Although the frequency of Ki-67^+^ cells among total CD8, Tn and Tcm was significantly decreased (**Fig. 2e, f**), the homeostatic proliferation of CD8 Tem was significantly augmented (**Fig. 2f**). Furthermore, the expression of PD-1 and Ki-67 showed a largely mutually-exclusive pattern among CD8 Tem (**Fig. 2g**). All these indicate that reduced PD-1 expression in the aged DJ-1 KO mice was not simply attributable to a general reduction in proliferation and activation under homeostatic conditions. Relevant cytokines, e.g., IFNg, increase among CD8 T cells during natural aging (Bandres et al. 2000). Therefore, we also analyzed IFNg production in CD8 T cells. Conforming to the other delayed immunoaging phenotypes, IFNg was decreased in CD8 T cells of aged DJ-KO versus WT mice following in vitro stimulation (**Fig. 2h**). Consistent with the observations in the index PD patient with DJ-1 deficiency, our results show that DJ-1 KO mice display reduced immunoaging in CD8 T-cell compartments.

T-cell-conditional deletion of the mitochondrial transcription factor A (*Tfam*), suppressing mitochondria DNA content, accelerates inflammaging (Desdin-Mico et al. 2020) and aging impairs mitochondrial homeostasis (Picca et al. 2018). To check whether our observation in cellular phenotypes can be extended to the organelle level we measured mitochondrial mass and membrane potential in different T cell subsets. As expected, both mitochondrial mass and membrane potential of different CD8 T-cell subsets (CD8 Tn, Tem and Tcm) were declined in aged WT mice versus young WT mice (**Fig. 2i-l, Extended Data Fig. 2c,d**). Notably, in both CD8 Tn and Tem from aged mice, the mitochondrial mass and membrane potential were significantly enhanced in DJ-1 KO versus WT mice (**Fig. 2i-l)**. Still similar to other CD8 subsets, CD8 Tcm mitochondria mass (p=0.07) and membrane potential (p=0.09) showed a tendency to increase in aged DJ-1 KO versus WT mice (**Extended Data Fig. 2c,d**). Consistent with other cellular data in young mice, significant difference was observed in neither mitochondrial mass nor membrane potential of different CD8 T-cell subsets between young DJ-1 KO and WT mice (**Fig.2i-l, Extended Data Fig. 2c,d)**. Thus, the concurring phenotypes between enhanced mitochondrial mass and functions in the CD8 subsets, especially in Tn and Tem, are in agreement with delayed immunoaging in cellular levels of aged DJ-1 KO versus WT mice in consideration of the observations in T cells devoid of *Tfam* (Desdin-Mico et al. 2020).

Although CD8 T cells are more susceptible to aging related changes (Czesnikiewicz-Guzik et al. 2008), we have also observed delayed immunoaging features in CD4 T cells of the index patent. Therefore, we also analyzed the murine CD4 T-cell compartments. As expected, similar effects were observed in CD4 T cells of aged DJ-1 KO mice (**Extended Data Fig. 3a-h**). To gain a more comprehensive picture, we performed transcriptomic analysis of CD4^+^CD25^-^ conventional T cells (Tconv) sorted from aged mice under homeostatic conditions. In accordance with the enhanced proportions of naïve T cells, our pathway enrichment analysis showed that several pathways involved in T-cell receptor signaling pathway and positive regulation of cell differentiation were significantly affected among the downregulated genes in Tconv of aged DJ-1 KO versus WT mice (**Extended Data Fig. 3l**). Notably, many memory T-cell or aging-related markers (Mogilenko et al. 2021) and/or memory T-cell development-related genes were also among the downregulated genes (**Extended Data Fig. 3l**). These results show that DJ-1 depletion also causes the delayed immunoaging in CD4 T-cell compartments.

**Figure 3.**
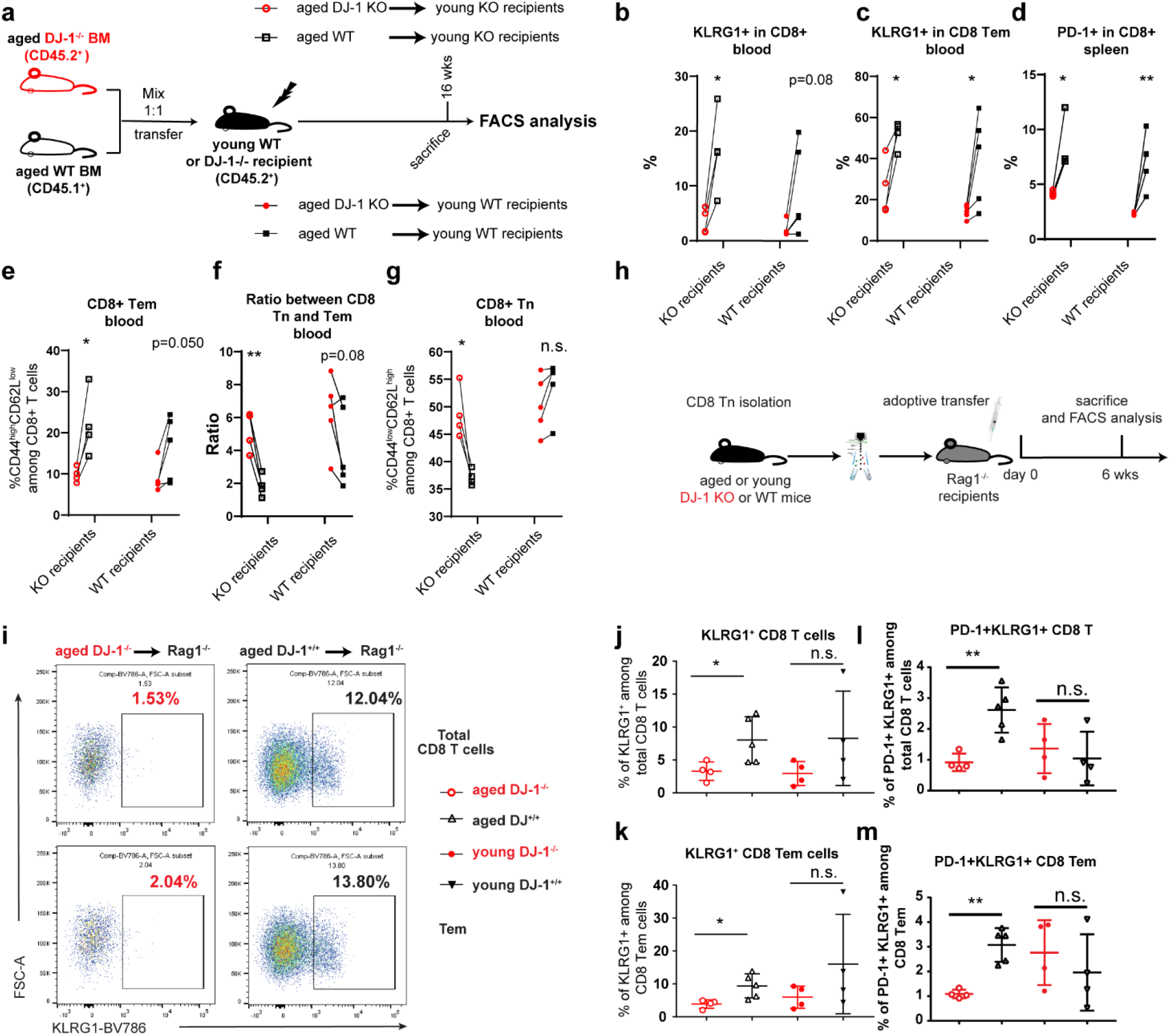
DJ-1 depletion inhibits the expression of key immunoaging markers KLRG1 and PD1 in a hematopoietic-and CD8-Tn-intrinsic manner. **a**, Schematic of the experimental setup of BM transplantation. A total of 10E6 of bone marrow cells from aged DJ-1 KO mice (CD45.2^+^) and WT mice (CD45.1^+^) (1:1 mix) were transferred into lethally-irradiated young DJ-1 KO or WT recipients (CD45.2^+^) by i.v. injection. Mice stably engrafted with donor cells were sacrificed for FACS analysis later. **b**,**d**, Percentages of KLRG1^+^(**b**) and PD-1^+^(**d**) CD8 T cells derived from aged DJ-1 KO and WT BM cells within young DJ-1 KO or WT recipients. **c**, Percentages of KLRG1^+^ among CD8 Tem derived from aged DJ-1 KO and WT BM cells within young DJ-1 KO or WT recipients. **e**,**g**, Percentages of CD8+ CD44^hi^CD62L^low^ (Tem) cells (**e**) and CD8+ CD44^low^CD62L^hi^ (Tn) cells (**g**) derived from aged DJ-1 KO and WT BM cells within young DJ-1 KO or WT recipients. **f**, Ratios between CD8 Tn and Tem cells developed from two types of BM cells within young DJ-1 KO or WT recipients. **h**, Schematic of the experimental setup of CD8 Tn adoptive transfer. 2.5E5 naïve CD8 T cells isolated from young or aged DJ-1 KO and WT littermates were injected into Rag-1 deficient mice by i.v. injection. 6 weeks later, mice were sacrificed for FACS analysis. **i**, Representative flow cytometry data of KLRG1^+^ population among total CD8 T (upper) and CD8 Tem cells (lower) following adoptive transfer of CD8 Tn from aged mice. **j, l**, Percentage of KLRG1^+^ (**j**) and KLRG1^+^PD-1^+^ (**l**) population among total CD8 T cells. **k, m**, Percentage of KLRG1^+^ (**k**) and KLRG1^+^PD-1^+^ (**m**) population among CD8 Tem cells. Results from BM transfer and adoptive transfer of CD8 Tn represent two independent experiments. Data are mean± s.d. The P-values are determined by a two-tailed paired (**b-g**) or non-paired (**j-m**) Student’s *t*-test. ns or unlabeled, not significant, *P<=0.05, **P<=0.01 and ***P<=0.001.

Since DJ-1 is a multi-functional protein ubiquitously expressed in different type of tissues and cells (Wilson 2011) and immunoaging involves many types of immune and non-immune cells (Nikolich-Žugich 2018), we next asked whether the delayed immunoaging in aged DJ-1 KO mice is the result of a hematopoietic-intrinsic or non-hematopoietic regulation. To this end, we generated mixed bone marrow (BM) chimeras by transferring BM cells mixed from young CD45.1 DJ-1 WT and CD45.2 DJ-1 KO donor mice into lethally-irradiated young WT recipients (**Extended Data Fig. 4a**). In this experiment, both DJ-1 KO and WT CD8 T cells developed under the same host WT environmental conditions. If the difference observed between that of KO and WT origins is consistent with the alteration in aged DJ-1 KO versus WT mice, it is attributable to hematopoietic intrinsic regulations. Interestingly, we indeed observed some hematopoietic-intrinsic phenotypes for the DJ-1-mediated immunoaging role. For instance, the expression of the critical immunoaging marker KLRG1 among blood CD8 T cells of DJ-1-KO origin was already significantly lower than that of DJ-1-WT-derived cells (**Extended Data Fig. 4b**) and also showed a trend to lessen in spleen (p=0.08, **Extended Data Fig. 4c**). This clearer phenotype in blood is in accordance with the notion that Tem are mainly distributed in non-lymphoid tissues (Masopust et al. 2001) and KLRG1 is mainly expressed among CD8 Tem. The exhaustion marker PD-1, which increases during aging, also showed a similar change as did KLRG1 (**Extended Data Fig. 4d**). The frequency of CD8 Tem was significantly lower in DJ-1-KO-derived cells (**Extended Data Fig. 4e,f**). Consistent with the data in aged DJ-1 KO mice, the ratios between CD8 Tn and Tem were already higher in the DJ-1-KO-derived cells than that from WT-origin cells (**Extended Data Fig. 4g,h**). These data indicate that DJ-1 exhibits a hematopoietic-intrinsic role for regulating not only the expression of immunoaging related markers, but also the proportion of CD8 Tem and the ratio between CD8 Tn and Tem.

However, the percentages of CD8 Tn did not show any significant difference between lymphocytes developed from young CD45.2 and CD45.1 BM cells in both spleen and blood (**Extended Data Fig. 4i,j**). Moreover, the percentages of CD8 Tcm derived from DJ-1 KO BM cells were higher than that from DJ-1*-*WT origin within young WT recipients (**Extended Data Fig. 4k,l**). Considering the observation in aged DJ-1 KO mice (**Extended Data Fig. 2a**), our results from young-donor-BM transplantation show that DJ-1 deficiency mediates accumulation of CD8 Tn and Tcm via either a hematopoietic-extrinsic or an aging-dependent manner or in combination of both. These mixture phenotypes, especially those non-intrinsic related observations, from the young-donor-BM chimeras urged us to investigate the potential effects of aging and non-hematopoietic aspects on the DJ-1-mediated immunaging.

To study the effects aforementioned, we generated mixed BM chimeras by transferring aged CD45.1 DJ-1 WT BM cells and CD45.2 DJ-1 KO BM cells into irradiated young WT or KO recipients (**Fig. 3a**). Due to the ethic limitation, we were unable to use aged mice as recipients. Similar to that in young-donor-BM chimeras (**Extended Data Fig. 4b-f**), the percentages of cells expressing the immunoaging related markers, e.g., KLRG1 (**Fig. 3b,c)** and PD-1 (**Fig. 3d**) as well as the frequency of CD8 Tem (**Fig. 3e**) were lower among CD8 cells reconstituted from aged DJ-1 KO versus WT BM donors, independent from the types of recipients, again supporting a hematopoietic-intrinsic mechanism. The ratios between CD8 Tn and Tem developed from aged DJ-1 KO versus WT BM donors were significantly higher in young KO recipients or with a tendency to increase also in young WT recipients (p=0.08, **Fig. 3f**). These consistent data from both young- and aged-donor-BM chimeras together suggests that DJ-1 exhibits an aging-BM-independent, but hematopoietic-intrinsic role for regulating the expression of KLRG1 and PD-1 among CD8 T cells, the frequency of CD8 Tem as well as the ratio between CD8 Tn and Tem.

Interestingly, the mixed BM chimeras did not show a significant difference in the frequency of CD8 Tn developed from aged DJ-1 KO versus WT BM donors as long as the recipients were WT mice, no matter from young (**Extended Data Fig. 4i,j)** or aged donors (**Fig. 3g**). Notably, consistent with the observation in aged DJ-1 KO versus WT mice, following reconstitution within young DJ-1-KO, but not -WT recipients, the proportion of blood CD8 Tn developed from aged DJ-1 KO versus WT BM cells was already significantly higher (**Fig. 3g**). At the same time, no significant change was observed in the frequency of CD8 single positive cells in thymus among aged DJ-1-KO versus -WT origins, within either type of young DJ-1 KO or WT recipients (**Extended Data Fig. 5a**), ruling out an abnormal thymic development of matured CD8 T cells. These results suggest that the enhanced frequency of CD8 Tn in aged DJ-1 KO versus WT mice requires the involvement of DJ-1-deficient non-hematopoietic cells. Opposed to the observations in aged DJ-1 KO versus WT mice, the percentages of CD8 Tcm of aged DJ-1-KO versus -WT origin were higher within both types of young recipients (**Extended Data Fig. 5b**,**c**). The results about CD8 Tcm percentages were consistent between young-and aged-BM models, regardless of within WT or KO recipients. These together essentially suggest the involvement of DJ-1-deficient aging microenvironmental factors in reduced accumulation of CD8 Tcm within aged DJ-1 KO mice. In short, our data firmly demonstrate that DJ-1 depletion enhances the ratio between CD8 Tn and Tem and inhibits the expression of immunoaging related markers in a hematopoietic-intrinsic but aging-BM-independent way, while regulating the CD8 Tn and Tcm accumulation in a more complicated manner.

The immunoaging process in T-cell compartments involves many types of cells (Nikolich-Žugich 2018) and BM-derived cells include far more than T cells. Previous data has shown that the homeostatic proliferation of CD8 Tn is enhanced during natural aging (Rudd et al. 2011) and the homeostatic proliferation of CD8 Tn drives the differentiation into CD8 Tem in Rag1^−/−^ lymphopenic mice (Cho et al. 2000). In aged DJ-1 KO versus WT mice, the homeostatic proliferation of CD8 Tn was reduced (**Fig. 2f**). Therefore, we hypothesized that this reduced homeostatic proliferation of CD8 Tn might impede or dysregulate the development into Tem cells and consequently the aging process of T cells. To test whether DJ-1 depletion has a T-cell-intrinsic effect on the transition from CD8 Tn into Tem subsets, we performed adoptive transfer of CD8 Tn sorted from young or aged DJ-1 KO or WT mice into young Rag1^−/−^ recipients, where the development of CD8 Tem is radically accelerated under lymphopenia (Cho et al. 2000) (**Fig. 3h**). Remarkably, the essential murine immunoaging marker KLRG1 was significantly lower in total CD8 T cells developed from aged DJ-1 KO vs WT donor cells (**Fig. 3i,j**). This phenomenon in total CD8 T cells was mainly due to the fact that from aged donor cells, CD8 T cells mainly consisted of CD8 Tem in this model (**Extended Data Fig. 6a-c**), where KLRG1 was significantly reduced in KO vs WT-originated cells (**Fig. 3k**). Furthermore, the frequency of the double-positive cells expressing both KLRG1 and PD1 among total CD8 T cells and CD8 Tem was significantly lower in aged DJ-1 KO CD8-Tn-originated cells (**Fig. 3l,m**). In line with the reduced expression of key immunoaging markers, the expression of the T-cell activation marker CD69 was significantly increased in total CD8 T cells and CD8 Tem derived from CD8 Tn of aged DJ-1-KO versus -WT origins (**Extended Data Fig. 6d,e**). Nevertheless, the frequency of CD8 Tn, Tem and Tcm in the *Rag1*-null recipients was not different in both comparisons of CD8 T cells from young DJ-1 KO versus -WT origins as well as that from aged DJ-1-KO versus -WT origins (**Extended Data Fig. 6a-c**), indicating the existence of CD8-Tn-extrinsic factors and/or requiring aging-microenvironmental factors. Nevertheless, the observed CD8-Tn-intrinsic role for DJ-1 in regulating key immunoaging related hallmarks in CD8 T cells, at least already partially, contribute to the delayed immunoaging phenotype in DJ-1 KO mice.

Our findings reveal an unexpected causal link between deficiency in a key monogenic PD gene PARK7/DJ-1 and delayed immunoaging in the T cell compartments. The data consistency between both human and mice with DJ-1 deficiency support a highly potent strategy to interfere with immunoaging for various complex diseases and infectious diseases, including COVID-19, where immunoaging is believed to be crucial (Koff and Williams 2020). Understanding the detailed molecular mechanisms through which DJ-1 regulates immunoaging still requires further investigation using cell-type-specific conditional DJ-1 KO mice. Our work also offers a unique animal model with delayed immunoaging phenotypes to allow researchers to explore the potential roles of a relatively juvenile immune system in the immune and aging associated diseases.

## Materials and Methods

Methods, list of materials used in this work and any related references are provided in Supplementary Information.

## Supplementary Information

In the initial submission, it is attached in the end of the pdf file. It will be available in the online version of the paper.

## Author contributions

N.Z. designed and performed major parts of the mouse-related experiments. N.Z. analyzed mouse-related data. C.C. performed and analyzed all the human and parts of mouse-related experiments. S.C. and A.B. performed parts of mice related experiments. D.C. and C.L. supervised parts of mouse-related experiments. H.K., J.B. A.W., D.B., R.K., R.B. and M.O. provided insights and supervised parts of the experiments. N.Z. and C.C. drafted the manuscript. R.K., M.O. and R.B. revised the manuscript. F.H. conceived and directed the project and revised the manuscript.

## Acknowledgements

We thank Annegrät Daujeumont, Nassima Ouzren, Caroline Davril and Julie Andre for their expert technical support. The group of F.H. was partially supported by Luxembourg National Research Fund (FNR) CORE programme grant (CORE/14/BM/8231540/GeDES), FNR AFR-RIKEN bilateral programme (TregBAR, F.H. and M.O.), PRIDE programme grants (PRIDE/11012546/NEXTIMMUNE and PRIDE/10907093/CRITICS). N.Z. and C.C. were supported through the PhD fellowship programme via PHD-2015-1/9989160 and PRIDE/2015/10907093, respectively. The work was also partially supported through intramural funding of LIH and LCSB through Ministry of Higher Education and Research (MESR) of Luxembourg.

## Extended Data Figures

**Extended Data Figure 1.**
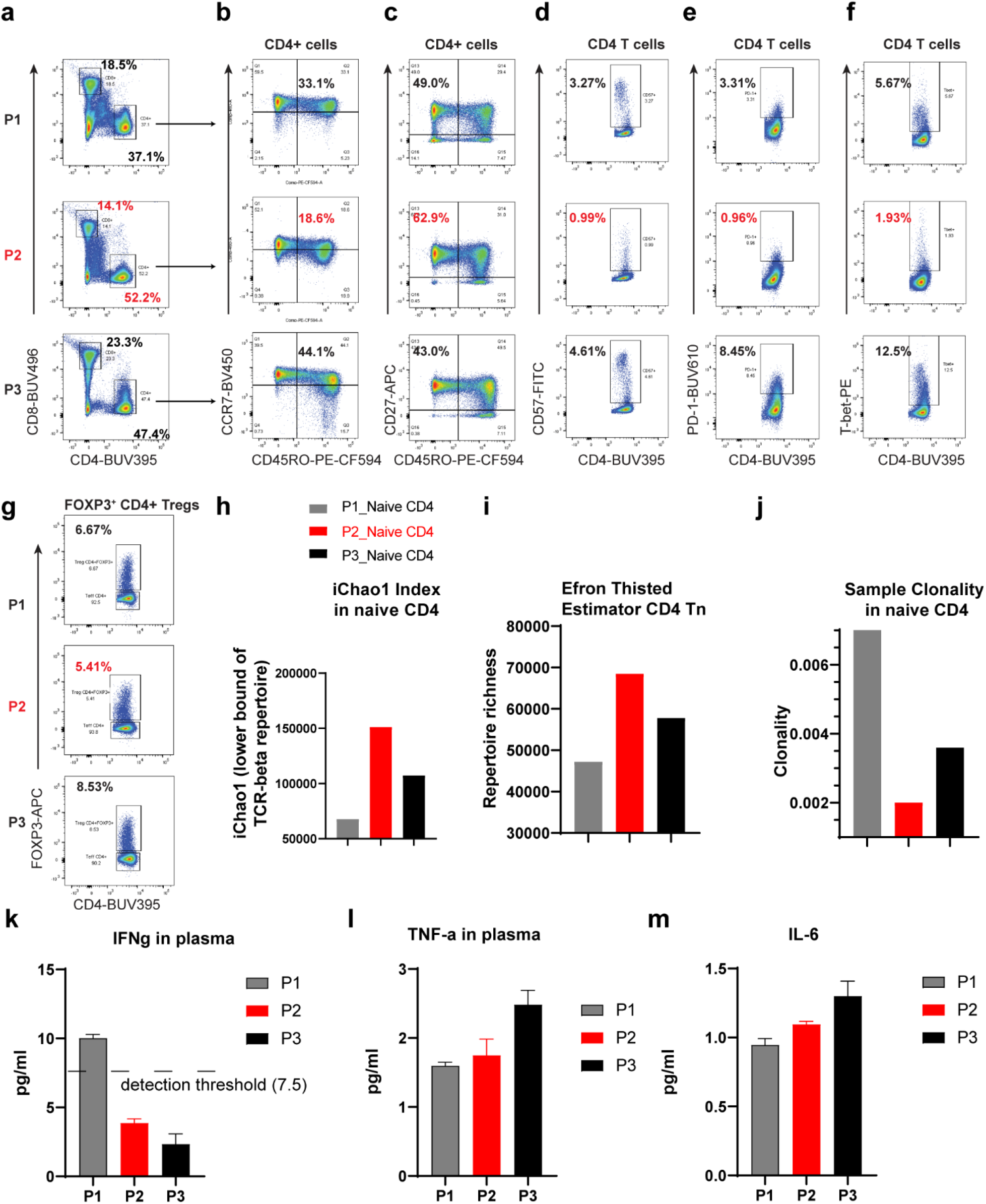
The DJ-1-devoid index patient showed delayed immunoaging features also in CD4 T cells. **a**, Percentages of total CD4 and CD8 T cells among the living lymphocyte singlets of peripheral blood of three participates [P1 (heterozygous mutation), P2 (homozygous mutation) and P3 (heterozygous)]. **b, c**, Coexpression of CCR7 and CD45RO (**b**), CD27 and CD45RO (**c**) on peripheral blood CD4 T cells of three participants. **d, e, f**, Expression of CD57 (**d**), PD-1 (**e**), and T-bet (**f**) on peripheral blood CD4 T cells of three participants. **g**, Frequency of FOXP3+CD4+ Tregs among total CD4 T cells in the peripheral blood from three participants. **h, i**, Comparison of the lower bound of TCR-beta repertoire (**h**) and richness (**i**) of sorted naïve CD4 T cells of three participants. **j**, The sample clonality index of TCR repertoire of naïve CD4 T cells of three participants. **k, l, m**, Cytokine measurement of IFNγ (**k**), TNF-α (**l**) and IL-6 (**m**) in the plasma of three participants. The other five tested cytokines were either undetectable or below fit curve. For cytokine measurement, data are mean± s.d.

**Extended Data Figure 2.**
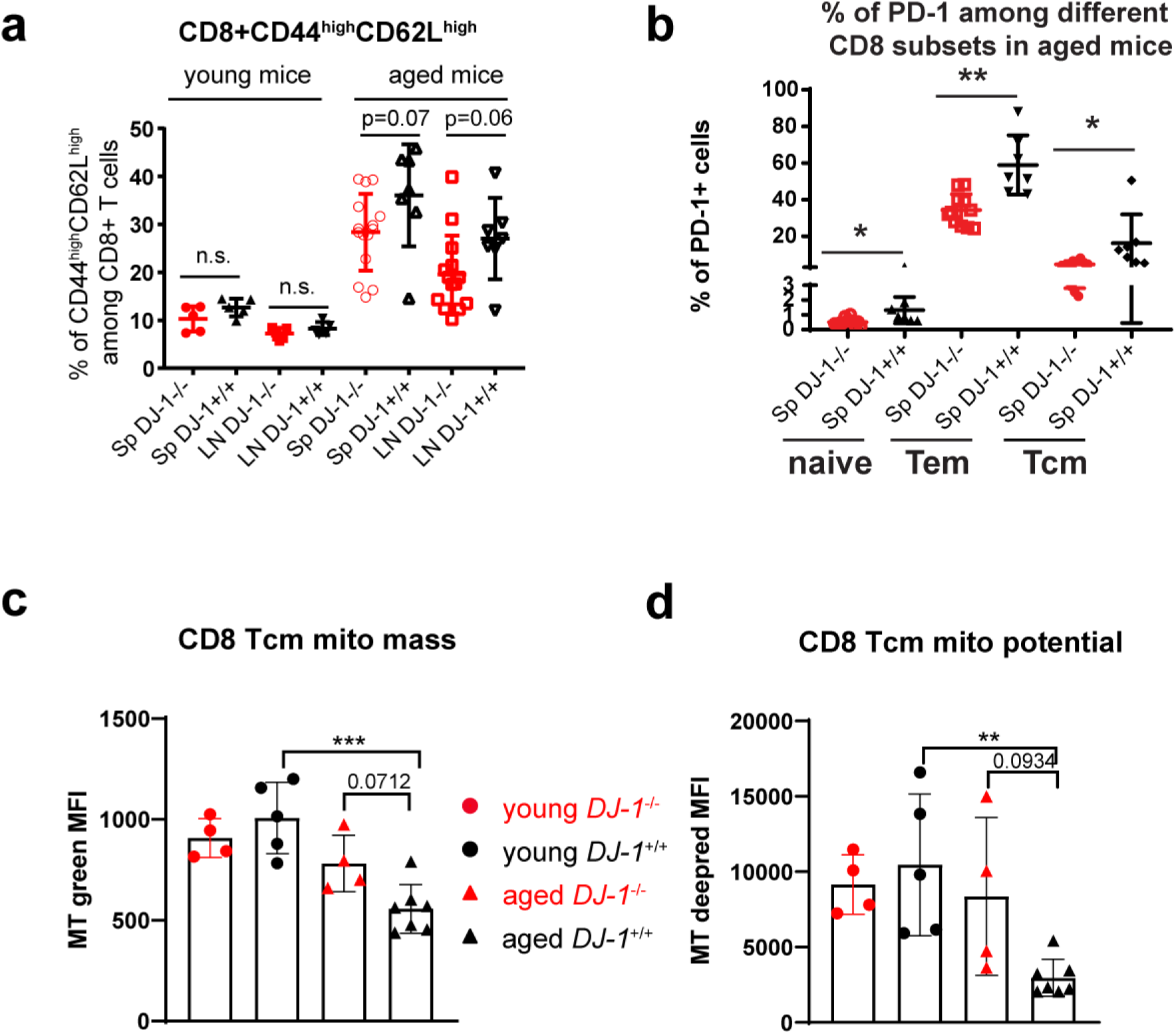
Extended characterization of delayed immunoaging in CD8 T-cell compartments. **a**, Percentages of CD44^high^ CD62L^high^ cells (Tcm) among total CD8 T cells of spleen and pLNs from young and aged DJ-1 KO and WT littermates. **b**, Percentage of PD-1^+^ cells among splenic CD8 Tn, Tem and Tcm subsets in the aged DJ-1 KO and age-and gender-matched WT mice. **c, d**, Comparison of CD8 Tcm mitochondrial (mito) mass (**c**) and mitochondrial potential (**d**) of young and aged DJ-1 KO and WT mice. Results represent at least four (**a, b**) and three (**c, d**) independent experiments. Data are mean± s.d.. The P-values are determined by a two-tailed non-paired Student’s *t*-test. n.s. or unlabeled, not significant, *P<=0.05, **P<=0.01 and ***P<=0.001.

**Extended Data Figure 3.**
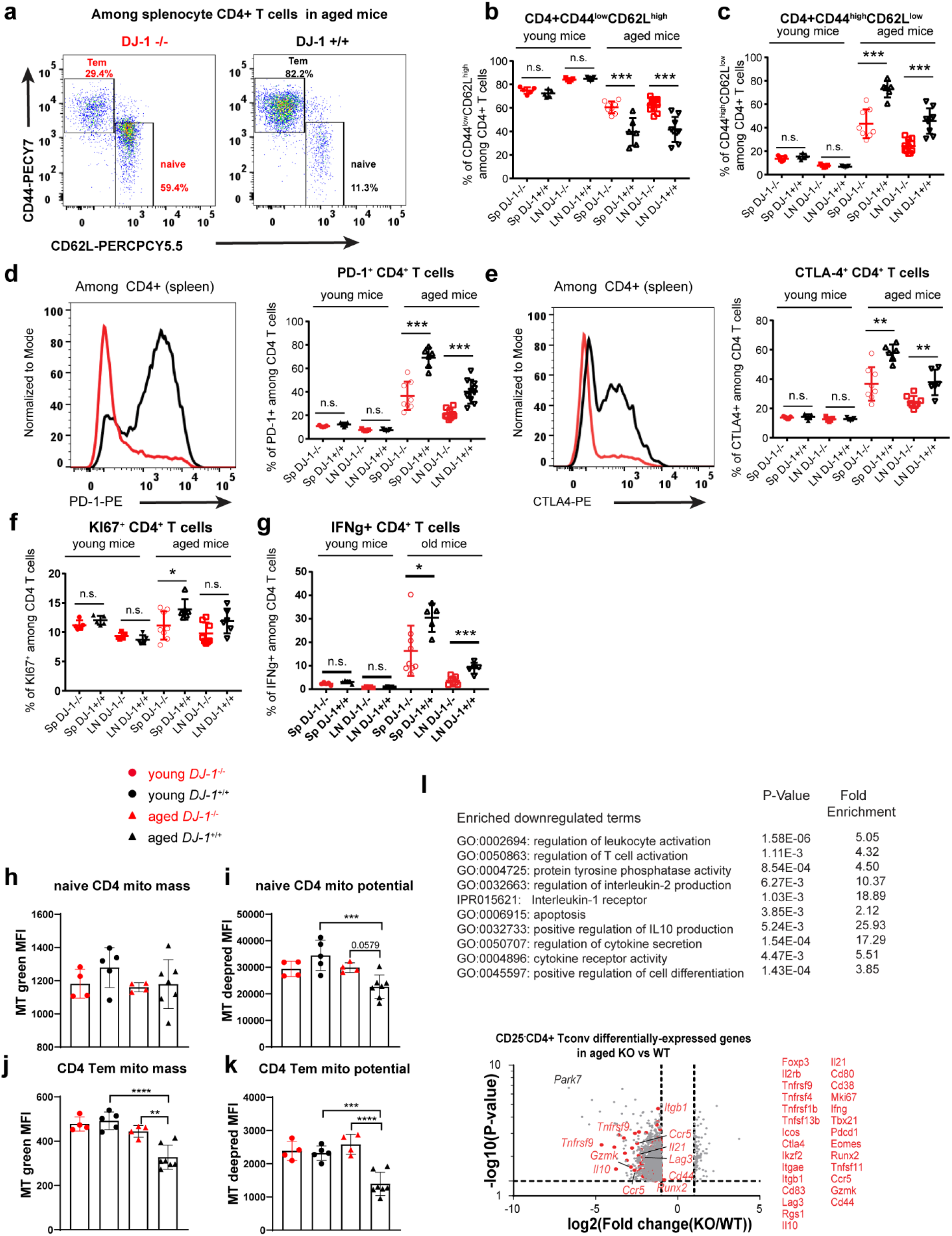
DJ-1 depletion renders the aged mice younger CD4 T cells. **a**, Representative flow-cytometry plots of CD44 and CD62L expression on total CD4 T cells of aged DJ-1 KO and age and gender-matched WT mice. **b, c**, Percentages of CD44^low^ CD62L^high^ (Tn) (**b**) and CD44^high^ CD62L^low^ (Tem) (**c**) cells among total CD4 T cells of spleen and pLNs from young and aged DJ-1 KO and WT littermates. **d**, Representative histogram overlay of PD-1 expression among total CD4 T cells in spleen of aged mice (left panel) and percentages of PD-1^+^ cells among total CD4 T cells (right panel). **e**, Representative histogram overlay of CTLA-4 expression among total CD4 T cells in spleen of aged mice (left panel) and percentages of CTLA-4^+^ cells among total CD4 T cells (right panel). **f**, Percentages of Ki-67^+^ cells among total CD4 T cells. **g**, IFNγ production in CD4 T cells of spleen and pLNs after in vitro stimulation using 50 ng/ml of PMA and 750 ng/ml of ionomycin for 5 h. **h, i**, Comparison of naive CD4 (Tn) mitochondrial (mito) mass (**h**) and mitochondrial (mito) potential (**i**) of young and aged DJ-1 KO and WT mice. **j, k**, Comparison of CD4 Tem mitochondrial mass (**j**) and mitochondrial potential (**k**) of young and aged DJ-1 KO and WT mice. **l**, The selected significantly enriched GO-terms and pathways among the downregulated genes in CD4 Tconv cells from aged DJ-1 KO mice versus the age-matched WT littermates from microarray analysis. **m**, Volcano plot shows both downregulated and upregulated differentially expressed genes in splenic CD4 T cells from three aged DJ-1 KO mice versus three age-matched WT littermates. SP and LN represents spleen and lymph nodes, respectively. Results represent at least four (**b-g**) and three (**h-k)** independent experiments. Data are mean± s.d. The P-values are determined by a two-tailed un-paired Student’s *t*-test. n.s. or unlabeled, not significant, *P<=0.05, **P<=0.01 and ***P<=0.001.

**Extended Data Figure 4.**
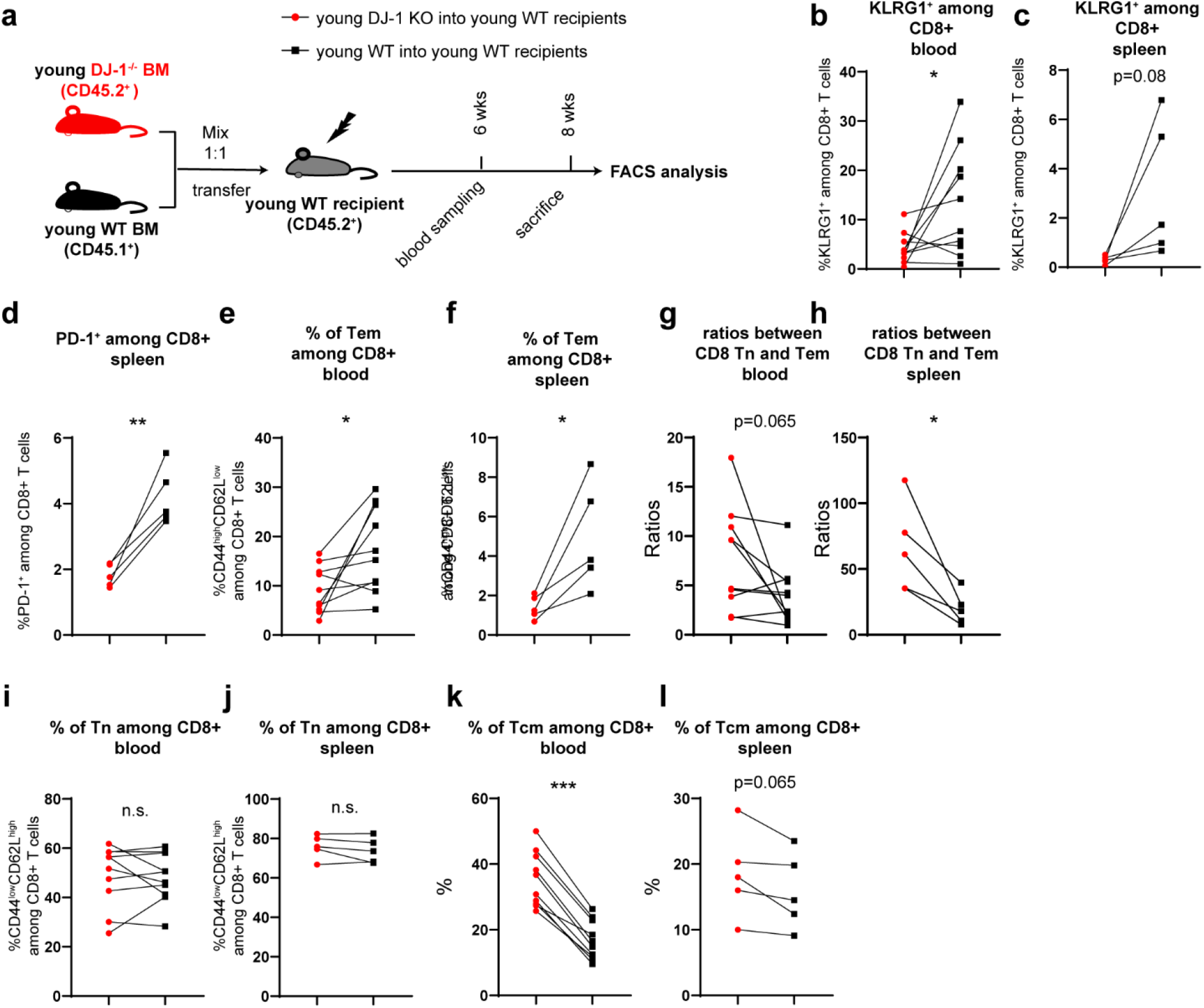
DJ-1 ablation regulates KLRG1 and PD-1 expression as well as the ratios between CD8 Tn and Tem in a hematopoietic-intrinsic manner but the accumulation of CD8 Tn and Tcm in a complicated manner. **a**, Schematic of the experimental setup of bone marrow transplantation. A total of 10E6 of bone marrow cells from young DJ-1 KO mice (CD45.2^+^) and WT mice (CD45.1^+^) (1:1 mix) were transferred into lethally-irradiated young WT recipients (CD45.2^+^) by i.v. injection. Mice stably engrafted with donor cells were sacrificed for FACS analysis later. **b, c**, Percentage of KLRG1^+^ CD8 T cells derived from young DJ-1 KO and WT donor BM cells in blood (**b**) and spleen (**c**) within young WT recipients. **d**, Percentage of PD-1^+^ cells among total CD8 T cells derived from young DJ-1 KO and WT BM cells in spleen within young WT recipients. **e, f**, Percentages of CD8 Tem derived from young DJ-1 KO and WT BM cells in blood (**e**) and spleen (**f**) within young WT recipients. **g, h**, Ratios between CD8 Tn and Tem cells developed from CD45.1 (WT) or CD45.2 (KO) BM cells in blood (**g**) and spleen (**h**) within young WT recipients. **i, j**, Percentages of CD8 Tn in blood (**i**) and spleen (**j**) derived from young DJ-1 KO and WT BM cells within young WT recipients. **k, l**, Percentage of CD8 Tcm among total CD8 T cells derived from young DJ-1 KO and WT BM cells in blood (**k**) and spleen (**l**) of young WT recipients. Results represent two independent experiments. The P-values are determined by a two-tailed paired Student’s *t*-test. n.s. or unlabeled, not significant, *P<=0.05, **P<=0.01 and ***P<=0.001.

**Extended Data Figure 5.**
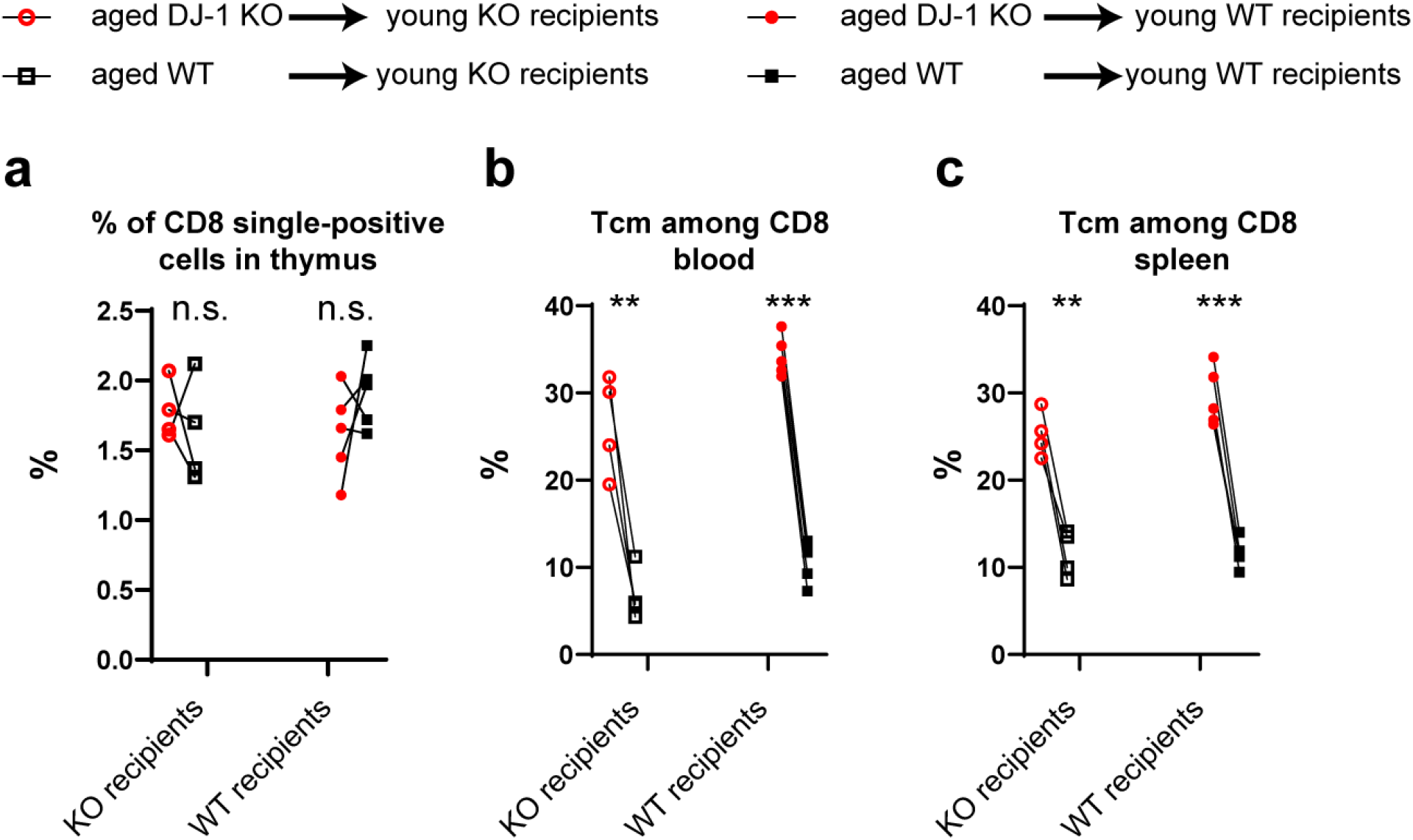
Extrinsic and intrinsic role for DJ-1 in regulating immunoaging related features in CD8 T-cell compartments. **a**, Percentages of CD8 single-positive cells among thymus originated from aged DJ-1 KO and WT BM cells within young DJ-1 KO or WT recipients following reconstitution. **b**,**c**, Percentages of CD8 CD44^high^CD62L^high^ (Tcm) cells in blood (**b**) and spleen (**c**) derived from aged DJ-1 KO and WT BM cells within young DJ-1 KO or WT recipients. Each symbol represent one mouse. Results represent two independent experiments. The P-values are determined by a two-tailed paired Student’s *t*-test. n.s. or unlabeled, not significant, *P<=0.05, **P<=0.01 and ***P<=0.001.

**Extended Data Figure 6.**
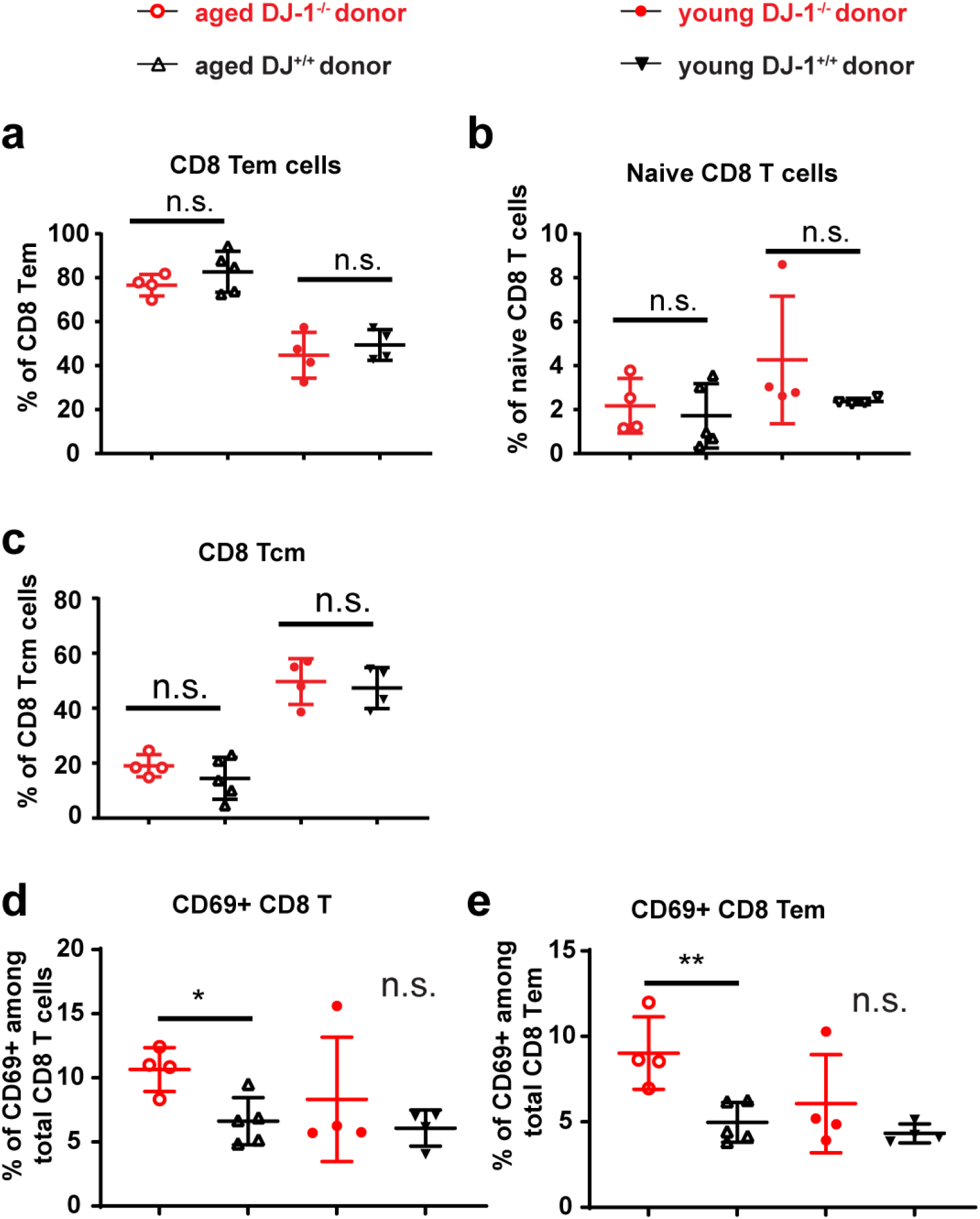
Extended characterization of CD8 T-cell compartments following adoptive transfer of CD8 Tn into *Rag1*-deficient mice. **a, b, c**, Percentages of splenic CD8 Tem (CD44^high^CD62L^low^) (**a**), Tn (CD44^low^CD62L^high^) (**b**) and Tcm (CD44^high^CD62L^high^) (**c**) cells. **d, e**, Percentage of CD69^+^ cells among total CD8 T cells (**d**) and CD8 Tem cells (**e**) in adoptive transfer experiment. Each symbol represent one mouse. Results represent two independent experiments. Data are mean± s.d. The P-values are determined by a two-tailed non-paired Student’s *t*-test. n.s. or unlabeled, not significant, *P<=0.05, **P<=0.01 and ***P<=0.001.

## Supplementary Information

### Materials and Methods

#### Animals

B6.129P2-Park7^Gt (XE726) Byg^/Mmucd mice were previously described^1^. The DJ-1 knockout mice line used in our lab has been generated by crossing the original line with B6N mice at least 10 generations. *Rag1*-/-mice with C57BL/6 (B6) background were purchased from the Jackson Laboratory and maintained in the specific pathogen free (SPF) animal facility of Luxembourg Institute of Health. CD45.1 mice with B6 background were purchased from Charles River Laboratory and maintained in the SPF animal facility of University of Luxembourg. The DJ-1^-/-^ (KO), DJ-1^+/-^, DJ-1^+/+^ (WT) mice used in our experiments were gender-and age-matched siblings generated from heterozygous DJ-1^+/-^ breeding pairs or trios. The DJ-1 line including both young and aged cohorts was also kept under the SPF conditions. Animal protocols have been approved either by animal welfare structure from LIH or animal experimentation ethics committee from University of Luxembourg, depending on where the experiments were performed. After institutional checking, each of the protocols was officially authorized by Luxembourg Ministry of Agriculture before starting the animal experiments.

#### Flow cytometry analysis for murine cells

The mice were sacrificed by dislocation of the neck. Spleens and peripheral lymph nodes (pLN) were collected and stored in the ice cold FACS buffer [Ca2+ and Mg2+ free PBS (Lonza, BE17-516F) with 2% inactivated FBS and 2mM EDTA, pH 8.0]. The pLNs here referred to a mixture of cervical, axillary and inguinal LNs. Spleens and lymph nodes were minced through a 70 µM cell cell strainer. Centrifuge the cell suspension at 350 g for 5 min at 4 °C. Discard the supernatant and add 3ml of 1x Red blood cell lysis buffer (BD, 555899) per mouse and incubate at room temperature for 7 min. Add 10 ml of FACS buffer to stop the reaction and centrifuge the cells and discard the supernatant. One million of cells per sample from spleen or pLNs were incubated in the FACS buffer [Ca2+ and Mg2+ free PBS (Lonza, BE17-516F) with 2% inactivated FBS and 2mM EDTA, pH 8.0] together with anti-mouse CD16/CD32 Fc blocker (BD Biosciences, 553141) at 4 °C for 15 min. Dead cells were distinguished using LIVE/DEAD^™^ Fixable Near-IR Dead Cell Stain Kit (Life Technologies, L34975). Then extracellular antibodies were added and incubated at 4 °C for 30 min protected from light. To detect intracellular nuclear proteins and transcriptional factors, cells were fixed and permeabilized using Foxp3/Transcription Factor Fixation kit (eBioscience, 00-5523-00) according to the manufacturer’s protocol. Samples were incubated with intracellular antibodies diluted in permeabilized buffer for 30 min at 4 °C. Cell acquisitions were performed on a BD LSRFortessa™ and data were analyzed using FlowJo software (v10, TreeStar, now part of BD). All the extracellular and intracellular antibodies are summarized in **Supplementary Table 1**. Of note, not necessarily all the abs listed there were analyzed in the same panel.

#### Mitotracker staining

Spleenocytes were isolated as previously described and stained for 30 min at 37 °C in complete RPMI medium containing anti-CD4 PE, anti-CD8 BUV737, anti-CD25 BUV395, anti-CD44 PE-Cy7, anti-CD62L PerCP-Cy5.5, 1:2000 DAPI, 1:100000 MitoTracker Green FM (ThermoFischer Scientific, M7514) and 1:10000 MitoTracker Deep Red (ThermoFischer Scientific, M22426). The cells were washed in cold FACS buffer and immediately acquired on a BD LSRFortessa™. The antibodies detecting surface markers for different subsets were provided in the **Supplementary Table 1**.

#### Intracellular cytokine quantification

For intracellular cytokine measurement, cells (2E5) from spleen or pLNs were stimulated by 50 ng/ml PMA (Phorbol 12-myristate 13-acetate, Sigma-Aldrich, P8139) and 750 ng/ml ionomycin (Sigma-Aldrich, I0634) in the presence of Golgiplug (BD Biosciences, 555029) and Golgistop (BD Biosciences, 554724) for 5 h in 96-well plates. Following cell surface staining, cells were fixed and permeabilized with Cytofix/Cytoperm buffer (BD Biosciences, 554714). The cytokine antibodies diluted in permeabilized buffer were added and incubated at 4 °C for 30 min protected from light. Cells were acquired on a BD LSRFortessa™ and analyzed by Flowjo v10.

#### Bone marrow transplantation

Bone marrow (BM) cells pooled from the femurs and tibias of DJ-1^-/-^ mice and age-, gender-matched CD45.1 B6 mice were isolated and 1:1 mixed in 100ul of cold PBS solution (Ca2+ and Mg2+ free; Lonza, BE17-516F). Gender-matched DJ-1 KO or WT mice aged 8-12 weeks as recipients were lethally irradiated from a gamma source (RS2000 X-Ray Biological Irradiator from Rad Source Technologies, two doses of 450 rads with 3 h resting period between the two doses) and 6 h later received 10E6 mixed donor BM cells by intravenous injection. For BM chimeras generated from young donor mice, blood was sampled at six and eight weeks post transplantation. Spleen was analyzed at eight weeks after transplantation. For BM chimeras generated from aged donor mice, both blood and spleen were analyzed at four months post transplantation. Cells were acquired on a BD LSRFortessa™ followed by the analysis with Flowjo v10.

#### Adoptive transfer of naïve CD8+ T cells

CD90.2^+^ cells from spleen and pLNs of DJ-1^-/-^ mice and DJ-1^+/+^ littermates aged 8-12 weeks or ∼55 weeks were magnetically isolated and first enriched by using anti-mouse CD90.2 microbeads (Miltenyi Biotec, 130-049-101, also refer to **Supplementary Table 2**). Naïve CD8^+^ T cells (CD3^+^CD8^+^CD44^low^CD62L^hi^) were then sorted by FACS sorting (BD FACSAria^™^III sorter (The sorting antibody information was already provided in **Supplementary Table 1**). 2.5E5 purified naïve CD8^+^ T cells in 100 ul of PBS (Lonza, BE17-516F) were injected intravenously into 8-12 week-old *Rag-1*^-/-^ mice. Six weeks post adoptive transfer, CD8^+^ T cells from spleens of the recipients were acquired by BD LSRFortessa^™^.

#### Microarray analysis of murine T cells

CD25^-^CD4^+^ Tconv cells from around 45-week-old DJ-1^-/-^ mice and DJ-1^+/+^ littermates were sorted using BD FACSAriaTM III sorter. The cell pellets were immediately lysated with RLT buffer supplemented with 1% beta-mercaptoethanol (Sigma-Aldrich, 63689) and frozen at -80 °C for further analysis. RNA was extracted by using the RNeasy Mini Spin Kit (Qiagen, 74104) and genome DNA was removed. Samples were analyzed via RNA 6000 Pico kit (Agilent, 50671513) by using an Agilent Bioanalyzer 2100, ensuring that only the samples have RIN higher than 8.5 were further used for microarray measurement.

RNA samples were analysed with the Affymetrix mouse Gene 2.0 ST Array at EMBL Genomics core facilities (Heidelberg). The microrray data analysis was performed in the same way as described in our previous work^2^. To ease the reading of this work, we described the major filtering steps here again. The expression signal at the exon level was summarized by the Affymetrix PLIER algorithm with DABG and PM-GCBG options by means of the sketch-quantile normalization approach (Affymetrix Expression Console v1.4). The corresponding probesets were considered differentially expressed if they passed the following combinatory filters^3^: (a) whether the change folds were >= 2 between the means of DJ-1 KO and WT Tconv; (b) whether the P-value, resulting from a two-tailed Student t-test, was <=0.05; (c) whether the cross-hyb type of the probeset was equal to 1; (d) whether the probeset with the highest expression level was higher than 100 (with the median value of ∼90 for each our sample); (e) for a given group (e.g. WT) with the higher mean intensity value of the probeset, whether the probeset in all the replicates of the given group was detected as ‘present’ according to the default setting of the Affymetrix Expression Console. DAVID v6.7 was used to perform functional enrichment analysis for the lists of differentially up-or down-regulated genes^4^. DJ-1-deficient Tconv cells exhibited much larger fractions of downregulated genes (416) compared with those of upregulated genes (53). This is why we focused on the analysis of the downregulated genes in this work.

#### PBMC isolation and flow cytometry analysis of the DJ-1-mutant patient family

We complied with all the relevant ethic regulations and Luxembourg CNER (Comité National d’Ethique de Recherche) has approved the PD patients related study. The family carrying the c.192G>C mutation in the DJ-1 gene has been already described elsewhere^5^ and written informed consent for all participating individuals was obtained. The index patient (56 years) carries the homozygous c.192G>C mutation and has been affected by PD for 22 years. The general disease progression over time is benign with a retained good response to levodopa therapy. The patient was currently presenting with a bilateral akinetic-rigid syndrome more pronounced on the left side with some postural instability and variable gait problems not interfering with his autonomy and corresponding to a Hoehn&Yahr stage 3. The two unaffected siblings (60 and 63 years) are both heterozygous carriers of the c.192G>C mutation and were devoid of any clinical sign of PD at the recent neurological examination, as expected for this autosomal-recessively inherited condition. The experimental investigators were blind to the genotype of the three siblings until the flow cytometry analysis was complete. PBMCs were isolated from the patient’s blood by Ficoll gradient centrifugation using SepMate tubes (StemCell, 86450, also refer to **Supplementary Table 3**) and Lymphoprep (StemCell, 07801). Following three washing steps 1E6 isolated PBMCs per patient for each staining panel were blocked for 15 min using Fc blocking antibodies (BD, 564765), then stained for 30 min at 4 °C using brilliant stain buffer (BD, 563794) containing the antibodies against cell surface markers from **Supplementary Table 4**. The cells were washed with human FACS buffer (PBS + 2% FBS, PBS is also Ca2+ and Mg2+ free; of note, slightly different from the FASC buffer for mice cell staining) and fixed for 1 h using the True-Nuclear Transcription factor buffer set (BioLegend, 424401). Following fixation, intracellular markers were stained for 30 min at room temperature in permeabilization buffer. After three washing steps, the cells were re-suspended in FACS buffer and acquired on a BD Fortessa. The flow cytometry data were analyzed using FlowJo software (v10, Tree Star).

#### TCR repertoire sequencing and mRNA microarray of the DJ-1-mutant patient family

Fresh PBMCs were stained with anti-CD4 FITC (BD, 555346, also refer to **Supplementary Table 4**), anti-CD8 BV605 (BioLegend, 301040) and LIVE/DEAD^™^ Fixable Near-IR Dead Cell Stain Kit (ThermoFischer Scientific, L34976) for 30 min at 4 °C. Stained cells were washed with human FACS buffer (PBS + 2%FBS) and total CD8 T cells were sorted using the BD Aria III. The cell pellets were immediately lysated with RLT buffer supplemented with 1% beta-mercaptoethanol (Sigma-Aldrich, 63689) and frozen at -80 °C for further analysis. The same procedure was applied for both human and mice cells to extract RNA (refer to the section above). The RIN values were analyzed by RNA 6000 Pico-kit (Agilent, 50671513) by using an Agilent Bioanalyzer 2100 and all the three samples had a RIN value of 10. Human RNA samples were analysed with the Affymetrix Human Gene 2.0 ST Array at EMBL Genomics core facilities (Heidelberg). A very similar procedure (refer to the section above) was applied to analyze both human and murine microarray datasets, with specific consideration of there were only three samples from human samples. The expression signal at the exon level was summarized by the Affymetrix PLIER algorithm with DABG and PM-GCBG options by means of the sketch-quantile normalization approach (Affymetrix Expression Console v1.4). The corresponding probesets were considered for further analysis if they passed the combinatory filters below: (a) the change folds were >= 2 between both the two comparisons P2 versus P1 and P2 versus P3; (b) the cross-hyb type of the probeset was equal to 1; (d) the probeset with the highest expression level was higher than 100; (e) if the expression of the given probeset was higher in the patient P2, the probeset in P2 had to be detected as ‘present’ according to the default setting of the Affymetrix Expression Console; if the expression of the given probeset was lower in the patient P2, the probeset in both P1 and P3 had to be detected as ‘present’.

PMBCs were cryopreserved in liquid nitrogen in aliquots of 5E6 cells in 1 ml (90% FBS + 10% DMSO). The thawed PBMCs were washed twice with warm (37 °C) supplemented complete IMDM (for the exact media components, refer to our previous work^2,6^) and recovered over night at 37 °C, 7.5% CO_2_. Antibodies used to stain for FACS sorting of naïve (CD3+CD45RO-CD45RA+) CD4+ or CD8+ T cells from patients’ PBMC are listed in **Supplementary Table 4**. Stained cells were washed with FACS buffer (PBS + 2%FBS) and naïve CD4 and CD8 T cells were sorted using the BD Aria III. DNA was extracted from the flash frozen pellets to perform TCR repertoire sequencing. Genomic DNA (gDNA) was extracted from the sorted naïve and memory CD4 and CD8 T cells using the QIAamp DNA Blood Mini Kit (Qiagen, 51104) following the manufacture’s instructions. The gDNA was eluted in 55 ul RNase-and DNase-free water to match the volume and concentration requirements for survey analysis of TCR beta repertoire sequencing by ImmunoSEQ (Adaptive Biotechnologies). All the analyses (TCR richness estimation and clonality) were performed using the online tool of ImmunoSEQ Analyzer 3.0. The lower bound of TCR repertoire of CD4 Tn and CD8 by applying nonparametric statistics using the iChao1 estimator. The richness of TCR repertoire of CD4 Tn and CD8 Tn by applying a nonparametric empirical Bayes estimation using the Efron Thisted estimator (extrapolation value is 120K from ImmunoSEQ Analyzer 3.0). The sample clonality of TCR repertoire of CD4 Tn and CD8 Tn (Clonality is equal to 1 – normalized Shannon’s Entropy).

#### Plasma cytokine measurement using the MSD platform

The concentration of the selected eight cytokines (IFNγ, IL-10, IL-17A, IL-1β, IL-4, IL-5, IL-6, TNFα) was measured in undiluted plasma samples of the three subjects using a multiplex MSD (Mesoscale Discoveries) U-plex Biomarker Group 1 (human) assay (MSD, K15067L) in the MSD MESO QuickPlex SQ 120 platform following the manufacturer’s instructions.

#### CMV ELISA

The CMV seropositivity was measured in the plasma samples of the DJ-mutant patients using the anti-Cytomegalovirus (CMV) IgG Human ELISA Kit (Abcam, ab108724) following the manufacturer’s instructions. The plasma was used at a 1:100 dilution.

## Data Availability

The microarray data from human CD8 T cells of the DJ-1 mutation carrier family and aged murine CD4 Tcon cells are deposited in Gene expression Omnibus (GEO) repository with the access code GSE173903 and GSE173904, respectively. To review the human CD8 dataset, go to https://www.ncbi.nlm.nih.gov/geo/query/acc.cgi?acc=GSE173903 and enter the token **avgryqwodlqnbqt** into the box. To review the mice dataset, go to https://www.ncbi.nlm.nih.gov/geo/query/acc.cgi?acc=GSE173904 and enter the token **utirsmiaxzyvxkn** into the box. Upon acceptance, we will make all those data including single-cell TCR repertoire sequencing data publically available.

## Supplementary Tables

**Supplementary Table 1.** List of mouse-related antibodies used in this work.

| Antibody | Clone | Company | Catalogue number | Dilution factor and application* |
| --- | --- | --- | --- | --- |
| CD16/CD32 | 2.4G2 | BD Biosciences | 553141 | 1:50 |
| CD3-BV421 | 145-2C11 | BD Biosciences | 562600 | 1:200 |
| CD3-APC | 17A2 | Biolegend | 100236 | 1:200 (sorting) |
| CD25-APC | PC61 | BD Biosciences | 557192 | 1:100 (sorting) |
| CD25-PE-Cy7 | PC61.5 | eBioscience | 25-0251-82 | 1:200 |
| CD25-BUV395 | PC61 | BD Biosciences | 564022 | 1:100(mitotracker) |
| CD44-PE-Cy7 | IM7 | BD Biosciences | 560569 | 1:200<br>(sorting) |
| CD4-BUV496 | GK1.5 | BD Biosciences | 564667 | 1:200 |
| CD4-FITC | RM4-5 | eBioscience | 11-0042-82 | 1:100 (sorting) |
| CD4-PE | RM4-5 | Biolegend | 100512 | 1:200 (mitotracker) |
| CD62L-PerCP-Cy5.5 | MEL-14 | BD Biosciences | 560513 | 1:200<br>(sorting) |
| CD8-BUV805 | 53-6.7 | BD Biosciences | 564920 | 1:200 |
| CD8-BV737 | 53-6.7 | BD Biosciences | 612759 | 1:200(mitotracker) |
| CD8-eFluor450 | 53-6.7 | eBioscience | 48-0081-82 | 1:100 (sorting) |
| CTLA4-PE | UC10-4B9 | eBioscience | 12-1522-83 | 1:100 |
| IFNγ-BV711 | XMG1.2 | BD Biosciences | 564336 | 1:100 |
| Ki-67-BV605 | 16A8 | BioLegend | 652413 | 1:200 |
| Ki-67-EF450 | SolA15 | eBioscience | 48-5698-82 | 1:100 |
| PD-1-BV711 | J43 | BD Biosciences | 744547 | 1:100 |
| KLRG1-PERCP-CY5.5 | 2F1/KLRG1 | Biolegend | 138417 | 1:100 |
| KLRG1-BV785 | 2F1/KLRG1 | Biolegend | 138429 | 1:100 |
| CD45.1-BV785 | A20 | Biolegend | 110743 | 1:200 |
| CD45.2-BV650 | 104 | Biolegend | 109835 | 1:200 |

\*, if not specified, the application is conventional FACS analysis; Specified application is mentioned in the bracket.

**Supplementary Table 2.** List of other materials used in mouse-related experiments of the study.

| **Materials** | **Company** | **Cat. Number** |
| --- | --- | --- |
| Agilent RNA 6000 Nano kit | Agilent | 5067-1511 |
| Agilent RNA 6000 Pico kit | Agilent | 5067-1513 |
| CD90.2 MicroBeads, mouse | Miltenyi Biotec | 130-049-101 |
| RPMI 1640 | Lonza | 12-167F |
| eBioscience™ Foxp3 / Transcription Factor Staining Buffer Set | eBioscience | 00-5523-00 |
| Fetal bovine serum (FBS) | Gibco | 10270106 |
| Fixation/Permeabilization Solution Kit | BD Biosciences | 554714 |
| Golgiplug | BD Biosciences | 555029 |
| Golgistop | BD Biosciences | 554724 |
| Ionomycin | Sigma-Aldrich | I0634 |
| Glutamax (100X) | Thermo Fisher Scientific | 35050061 |
| Sodium Pyruvate | Thermo Fisher Scientific | 11360070 |
| HEPES | Thermo Fisher Scientific | 15630080 |
| PBS | Lonza | BE17-516F |
| PMA | Sigma-Aldrich | P8139 |
| Red blood cell lysis buffer | BD | 555899 |
| RNeasy Mini Spin Kit | Qiagen | 74104 |
| DNase I | Qiagen | 79254 |
| MEM non-essential amino acids | Sigma-Aldrich | M7145 |
| 0.5 M EDTA | Sigma-Aldrich | E7889 |
| Penicillin+Streptomycin | Thermo Fisher Scientific | 15070-063 |
| Beta-Mercaptoethanol | Sigma-Aldrich | M7522 |

**Supplementary Table 3.** Materials or reagents used for human PBMC isolation and flow cytometry analysis.

| **Name** | **Company** | **Ref. number** |
| --- | --- | --- |
| SepMate tubes | StemCell | 86450 |
| Lymphoprep | StemCell | 07801 |
| True-Nuclear Transcription Factor Buffer Set | BioLegend | 424401 |
| Brilliant Stain Buffer | BD | 563794 |
| UltraComp eBeads | eBioscience | 01-2222-42 |

**Supplementary Table 4.** List of antibodies used for sorting or analysing human T cells.

| Protein | Color | Dilution | Company | Ref. number | Clone |
| --- | --- | --- | --- | --- | --- |
| **Analysis staining abs** |  |  |  |  |  |
| Fc Blocking Abs | / | 1:50 | BD | 564765 | / |
| CD4 | BUV395 | 1:40 | BD | 563550 | SK3 aka Leu3a |
| CD8 | BUV496 | 1:40 | BD | 564804 | RPA-T8 |
| CD27 | APC | 1:40 | BD | 561786 | M-T271 |
| CD28 | BUV785 | 1:40 | BioLegend | 302950 | CD28.2 |
| CD45RO | PE-CF594 | 1:40 | BD | 562299 | UCHL1 |
| CD57 | FITC | 1:40 | BD | 555619 | NK-1 |
| CD197 (CCR7) | Pacific Blue | 1:40 | BioLegend | 353210 | G043H7 |
| Eomes | PE-Cy7 | 1:20 | Thermo Fischer Scientific | 25-4877-42 | WD1928 |
| FOXP3 | APC | 1:20 | BioLegend | 320114 | 206D |
| Ki-67 | FITC | 1:20 | BD | 561165 | B56 |
| PD-1 | BV605 | 1:40 | BioLegend | 329924 | EH12.2H7 |
| T-bet | PE | 1:20 | BioLegend | 644810 | 4B10 |
| Live/Dead | APC-Cy7 | 1:500 | Thermo Fischer Scientific | L34976 | / |
| **Microarray analysis sorting panel** |  |  |  |  |  |
| CD4 | FITC | 1:20 | BD | 555346 | RPA-T4 |
| CD8 | BV605 | 1:20 | BioLegend | 301040 | RPA-T8 |
| Live/Dead | APC-Cy7 | 1:500 | Thermo Fischer Scientific | L34976 | / |
| **TCR repertoire analysis sorting panel** |  |  |  |  |  |
| Fc Blocking Abs | / | BD Bioscience | / | 564765 | 1:50 |
| CD3 | HIT3a | BD Bioscience | BV510 | 741822 | 1:100 |
| CD4 | RPA-T4 | BD Bioscience | FITC | 555346 | 1:100 |
| CD8 | RPA-T8 | Biolegend | BV605 | 301040 | 1:100 |
| CD45RA | HI100 | Biolegend | Pacific Blue | 304123 | 1:50 |
| CD45RO | UCHL1 | BD Bioscience | PE-CF594 | 562299 | 1:50 |

